# Studying the history of tumor evolution from single-cell sequencing data by exploring the space of binary matrices

**DOI:** 10.1101/2020.07.15.204081

**Authors:** Salem Malikić, Farid Rashidi Mehrabadi, Erfan Sadeqi Azer, Mohammad Haghir Ebrahimabadi, S. Cenk Sahinalp

## Abstract

Single-cell sequencing data has great potential in reconstructing the evolutionary history of tumors. Rapid advances in single-cell sequencing technology in the past decade were followed by the design of various computational methods for inferring trees of tumor evolution. Some of the earliest of these methods were based on the direct search in the space of trees. However, it can be shown that instead of this tree search strategy we can perform a search in the space of binary matrices and obtain the most likely tree directly from the most likely among the candidate binary matrices. The search in the space of binary matrices can be expressed as an instance of integer linear or constraint satisfaction programming and solved by some of the available solvers, which typically provide a guarantee of optimality of the reported solution. In this review, we first describe one convenient tree representation of tumor evolutionary history and present tree scoring model that is most commonly used in the available methods. We then provide proof showing that the most likely tree of tumor evolution can be obtained directly from the most likely matrix from the space of candidate binary matrices. Next, we provide integer linear programming formulation to search for such matrix and summarize the existing methods based on this formulation or its extensions. Lastly, we present one use-case which illustrates how binary matrices can be used as a basis for developing a fast deep learning method for inferring some topological properties of the most likely tree of tumor evolution.

## 1 Introduction

Cancer is an evolutionary disease characterized by successive rounds of mutation and selection. Deciphering the evolutionary history of a tumor represents an important problem in cancer studies and can help us in better understanding some of the key aspects of tumor initiation, progression, metastatic spread and many others.

Rapid advances in sequencing technologies in the past decade were followed by the development of various computational methods for studying tumor evolution (Schwartz and Schäffer, 2017; Kuipers et al., 2017). Most of the earliest methods were designed for traditional bulk sequencing data, which is obtained by sequencing a mixture of DNA originating from a large number of cells. Since bulk data does not contain information about the cell of the origin of a particular read, studying tumor evolution by the use of this data type is very challenging. For example, in many cases, it is impossible to unambiguously differentiate between several topologically very distinct trees of tumor evolution that describe a given bulk dataset equally well. We refer to (Kuipers et al., 2017) and (Malikic et al., 2019a) for a more detailed discussion of limitations of the use of bulk data in this context.

The second main group of methods for inferring tumor evolutionary history consists of methods designed for single-cell sequencing (SCS) data. Due to its high potential in studying tumor evolution, the advent of single-cell sequencing (SCS) has attracted a lot of attention in the field of tumor phylogenetics. In contrast to the bulk data, the high resolution data generated by sequencing individual tumor cells enables researchers to infer tumor evolutionary history at unprecedented detail. However, due to the elevated noise rates present in a typical SCS dataset, inferring trees of tumor evolution from SCS data is not straightforward and it necessitated the development of automated computational methods. In the past few years, we have been witnessing very high interest in the design of such methods and many novel and sophisticated methods were proposed. They are based on the use of various techniques including heuristics, Markov Chain Monte Carlo (MCMC) search, integer linear programming (ILP), constraint satisfaction programming (CSP) and others.

In this review, we first provide a brief description of the mutation tree of tumor evolution and binary genotype matrix obtained from SCS data. Next, we summarize the tree scoring scheme commonly used in the existing methods for finding the maximum likelihood tree of tumor evolution from SCS data. We then provide an overview of the maximum likelihood tree search strategy first proposed in SCITE (Jahn et al., 2016), one of the earliest methods for SCS data. Our main goal here is describing how the search in the space of trees performed in SCITE can be replaced by search in the space of conflict-free binary matrices (see below for the definition). We show how the search in the space of matrices can be expressed as an instance of ILP and routinely solved by the use of the available ILP solvers. One of the main advantages of such implementation in comparison to the methods based on MCMC search is the availability of some measure of the quality of reported solutions (ideally, a guarantee of optimality), which is provided by ILP solvers. The potential of tree inference by performing a search in the matrix space has already been recognized by several research groups and here we summarize the existing methods based on this strategy. Finally, we introduce a class of staircase matrices, which represent a subclass of conflict-free matrices and have recently been successfully exploited in the time-efficient deep learning-based approach for identifying the presence of divergent subclones in a given tumor (Sadeqi Azer et al., 2020a).

Note that here we focus on somatic single-nucleotide variants (SNVs) as genetic footprints of cancer evolution and any use of the term “mutation” hereafter refers to this type of genetic mutation. In addition, unless otherwise stated, we assume that commonly used infinite sites assumption (ISA) holds. This assumption states that each mutation is acquired exactly once during the course of tumor evolution and is never lost (i.e., it is inherited by all descendants of the cell where it occurred for the first time).

## 2 Mutation tree of tumor evolution

One of the most convenient ways of depicting the evolutionary history of a given tumor is by the use of mutation tree (Jahn et al., 2016). A mutation tree on mutations *M*_1_, *M*_2_, …, *M*_*m*_ is a rooted tree which consists of *m* + 1 nodes and each of its edges is labeled by exactly one of these *m* mutations (due to ISA each mutation is assigned to exactly one edge so we have a one-to-one correspondence between the tree edges and the mutations). Nodes of a given mutation tree can be labeled arbitrarily. Here, we use labeling with non-negative integers with root being labeled by 0. A simple example of mutation tree is shown in Figure 1. Note that in this figure one should focus only on green nodes, solid edges and their labels as the main constituents of the mutation tree. In the figure we also show nine single-cells (yellow nodes) sampled in some SCS experiment.

**Figure 1.**
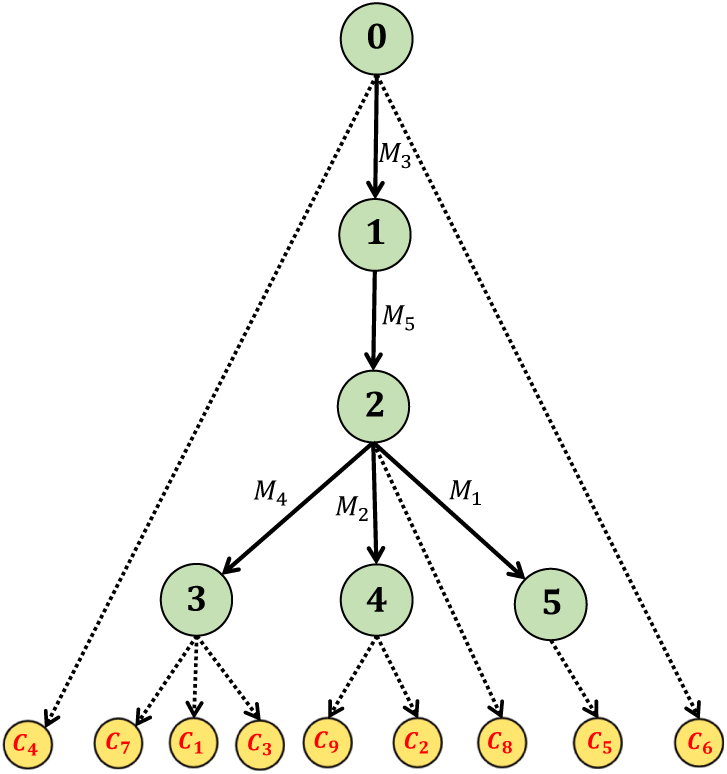
An example of mutation tree (green nodes, solid edges) from which nine single-cells *C*_1_, *C*_2_, …, *C*_9_ were sampled. Each single-cell is connected to its node of origin in the mutation tree via a dashed edge.

Each node *i* of the mutation tree can be associated with a binary genotype vector *G*_*i*_ = [*G*_*i*1_, *G*_*i*2_, …, *G*_*im*_]. We assume that the root node is mutation-free and define that *G*_0*p*_ = 0 for all *p*. For any other node *i, G*_*ip*_ = 1 if and only if mutation *M* occurs on the path from the root to the node *i*. Intuitively, *G*_0_, *G*_1_, …, *G*_*m*_ represent all possible genotypes present during the course of tumor evolution, where genotypes are defined on the set of *m* mutated loci. The first cell with genotype *G*_*i*_ ≠ *G*_0_ is born when some cell with genotype *G*_*j*_ acquires mutation *M*_*p*_, where node *j* is a parent of node *i* and *M*_*p*_ is the label of the edge connecting *j* and *i* (note that in this sentence we assume that each daughter cell differs from its parent cell by at most one mutation, but the assumption is made solely for simplifying the presented intuitive description of *G*_*i*_). Note also that, since we expect that each of the cells originates from one of the nodes of the tree, the true genotype of the cell is expected to match some of the node genotypes *G*_0_, *G*_1_, …, *G*_*m*_.

For a given mutation tree and two distinct mutations *M*_*p*_ and *Mq* we define that *M*_*p*_ is an ancestor of *M*_*q*_ if and only if the edge labeled by *M*_*p*_ is contained within the path starting at the root of the tree and ending at the edge labeled by *M*_*q*_ (including this edge). If neither *M*_*p*_ is an ancestor of *M*_*q*_ nor *M*_*q*_ is an ancestor of *M*_*p*_ than we say that *M*_*p*_ and *M*_*q*_ belong to different branches of the tree.

## 3 Inference of tumor evolutionary history from single-cell sequencing data

### 3.1 Binary genotype matrix obtained from single-cell sequencing data

Assume that we have performed a single-cell sequencing experiment where *n* single-cells, denote them as *C*_1_, *C*_2_, …*C*_*n*_, were sampled from the same patient. The true genotypes of these cells can be arranged in a single-cell binary genotype matrix *A* with the columns of *A* corresponding to mutations and the rows representing genotypes of single-cells. One example of such a matrix is shown in the Figure 2.

**Figure 2.**
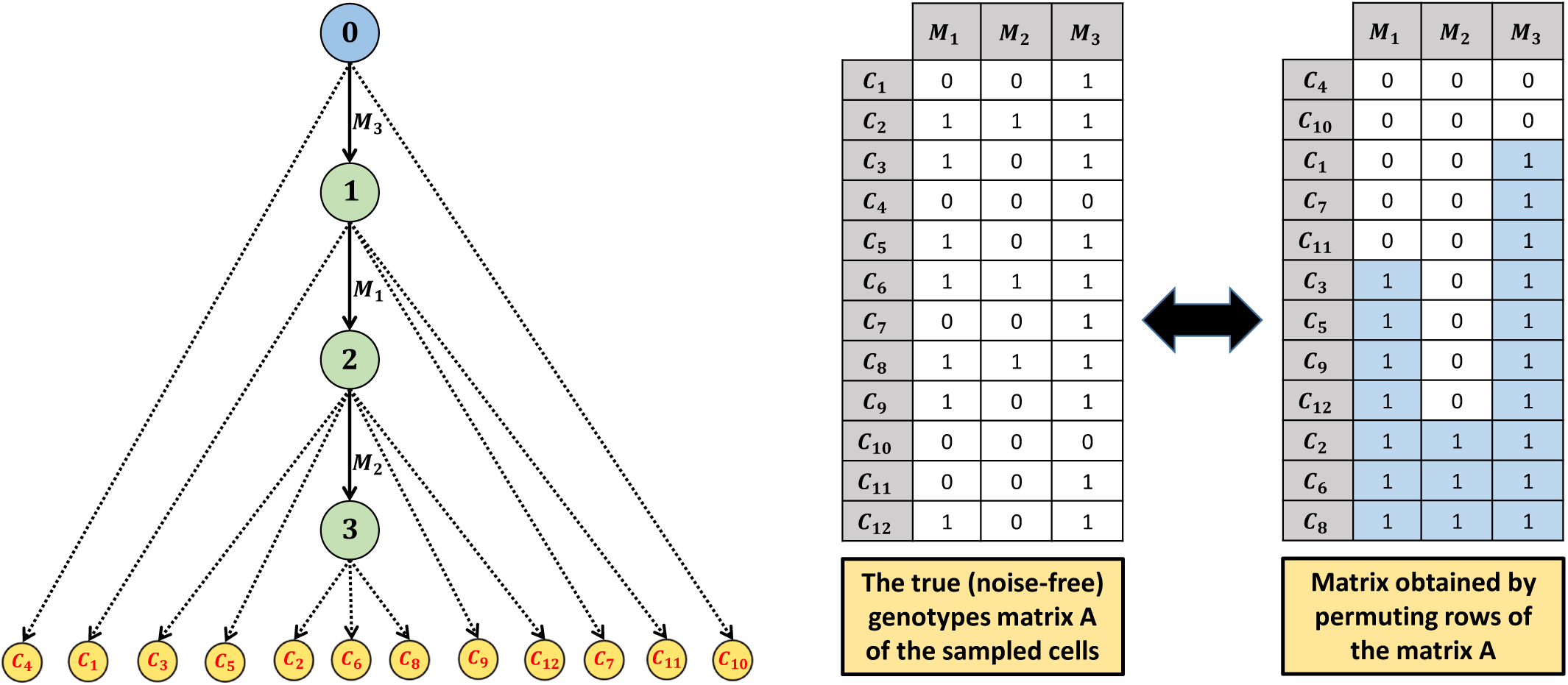
Mutation tree with linear topology from which twelve single-cells were sampled (left). True single-cell genotype matrix *A* of the sampled single-cells (middle). The matrix obtained by permuting rows of *A* in order to obtain a matrix *S* such that the inequalities *S*_*ij*_ ≤ *S*_(*i*+1)*j*_ from the definition of staircase matrix are satisfied (right).

Due to the noise present in SCS data, it is usually very unlikely to observe the matrix *A* directly from the raw sequencing data. Assume that, after processing the raw sequencing data and based on some mutation calling criteria, we found an evidence for *m* putative mutations that we denote as *M*_1_, *M*_2_, …, *M*_*m*_. Assume for simplicity, that for each single-cell and each mutation we make a binary decision about the presence or absence of the mutation in the cell (later we will also reference more general models where, instead of making a binary choice, variant and reference read counts at the mutation loci are considered in order to get a confidence score for mutational presence). Then the output of the above steps can be represented as a binary matrix *D* consisting of *n* rows and *m* columns which respectively correspond to the sets of sequenced single-cells and putative mutations. The value of *D*_*ij*_ is set to 1 if our mutation calling procedure predicted that mutation *M*_*j*_ is present in cell *C*_*i*_. Otherwise, we set *D*_*ij*_ = 0.

### 3.2 Scoring an arbitrary mutation tree

Given a matrix *D* introduced above, our goal is to find a mutation tree that best explains *D*. Here we describe how an arbitrary mutation tree is scored. Note that we focus on the trees defined on the set of mutations {*M*_1_, *M*_2_, …, *M*_*m*_ } as the only candidate solutions.

As mentioned earlier, for a given mutation tree, we expect that each of the sequenced single-cells originates from one of the tree nodes. The most commonly used way of defining a score of a particular tree *T* is the following: (i) for a given row *i* in *D* (i.e., cell *C*_*i*_), compute the similarity score of the row vector [*D*_*i*1_, *D*_*i*2_, …, *D*_*im*_] to the genotype of each of the tree nodes (ii) Among all of the computed scores, find the largest one and set it as the score of the cell *C*_*i*_. Per this solution, the true genotype of the cell *C*_*i*_ then equals to the genotype associated with the node for which the largest score is attained. (iii) Repeat the previous two steps for all rows and compute the score of *T* by combining the scores of individual rows.

Two important questions that remain are: (i) definition of the similarity score between rows of *D* and genotypes of tree nodes and (ii) the algorithm for searching the space of all candidate trees.

For defining the similarity score, first note that in the above we are associating a binary vector *D*_*i*_ = [*D*_*i*1_, *D*_*i*2_, …, *D*_*im*_] with a binary vector *E*_*i*_ = [*E*_*i*1_, *E*_*i*2_, …, *E*_*im*_], where *E*_*i*_ ∈ {*G*_0_, *G*_1_, …, *G*_*m*_ }. Pairs (*i, j*) such that (*D*_*ij*_, *E*_*ij*_) = (1, 0) imply a false positive mutation call in *D* for cell *C*_*i*_ and mutation *M*_*j*_. Similarly, a pair (*D*_*ij*_, *E*_*ij*_) = (0, 1) implies a false negative mutation call. It is well known that false positive and false negative noise rates in SCS data are usually very different. While false positives are typically rare (at rate less than *<* 0.01), for most of the available datasets false negative rate is estimated to be between 0.1 and 0.3. Therefore defining the similarity score for *D*_*i*_ and arbitrary *G*_*v*_ via naive counting of the number of coordinates at which these two vectors are equal would not properly discriminate between the two noise types. Instead, what is commonly adopted is the use of probability of observing genotype *D*_*i*_ given that the true genotype is *G*_*v*_. Assuming that the mutated sites evolve independently and denoting the false positive and false negative noise rates of SCS data as *α* and *β*, respectively, this probability equals

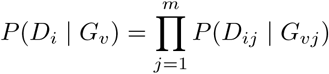

Where

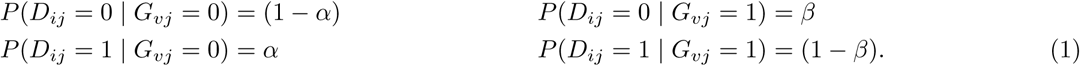

The genotype *E*_*i*_, defined above as the most similar to *D*_*i*_ among all node genotypes, is then given by the following formula

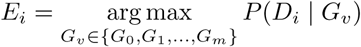

and the likelihood score of the tree *T*, denoted as *S*(*T*), is given as

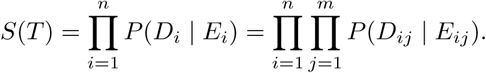

### 3.3 Tree inference by search in the space of mutation trees

The first approach for finding the maximum likelihood tree that we summarize here is a direct search in the space of mutation trees on *M*_1_, *M*_2_, …, *M*_*m*_, denoted below as 𝕋. This approach was first introduced in (Jahn et al., 2016) and starts with a random tree from 𝕋, which is set as a *current* tree. Given the current tree *T*, the new tree *T* ′ is proposed by the use of several types of moves in the tree space (e.g., swapping labels of two arbitrary edges of *T*). Based mostly on the scores *S*(*T*) and *S*(*T* ′) the proposed tree *T* ′ is either accepted and becomes the new current tree, or is rejected. The tree proposing step is repeated for a given number of iterations and the whole procedure can be restarted for a given number of repeats, both specified by a user. In the end, the best-found tree is returned as the most likely evolutionary history of the sequenced tumor.

### 3.4 Tree inference by search in the space of conflict-free matrices

We will now present an alternative approach for finding the most likely evolutionary history of a tumor that is based on the search in the space of *conflict-free* matrices. A conflict-free matrix is any binary matrix *X* such that for each triplet of rows *i, j, h* and each pair of columns *p* and *q*

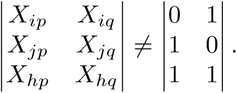

If there exists a triplet of rows and a pair of columns for which equality holds in the above, then we say that they form a *conflict* (Hujdurović et al., 2015). We denote the space of all conflict-free matrices of size *n* × *m* as 𝕄.

We will now first show that the maximum likelihood mutation tree can be obtained from the best scoring conflict-free matrix and then provide an exact ILP formulation for finding such matrix.

#### 3.4.1 Search for the maximum likelihood tumor evolutionary history in the space of mutation trees vs. the space of conflict-free matrices

Using the same notation as above, let *T* be an arbitrary mutation tree and let *E* denote the maximum likelihood matrix associated with it. It can be easily shown that matrix *E* is conflict-free. Namely, if *M*_*p*_ and *M*_*q*_ are two arbitrary mutations then either one of them is ancestor of the other or they belong to different branches of the tree. If *M*_*p*_ is ancestor of *M*_*q*_ then, due to the infinite sites assumption, any cell harboring *M*_*q*_ also harbors *M*_*p*_, implying that (*E*_*ip*_, *E*_*iq*_) ≠ (0, 1) for each row *i*. Similarly, if *M*_*q*_ occurs before *M*_*p*_ then (*E*_*ip*_, *E*_*iq*_) ≠ (1, 0) for each row *i*. On the other hand, if *M*_*p*_ and *M*_*q*_ belong to different branches of the tree then obviously (*E*_*ip*_, *E*_*iq*_) ≠ (1, 1) for each row *i*. Hence, regardless of the relative placement of mutations *M*_*p*_ and *M*_*q*_ in *T*, at least one of the triplets (1, 0), (0, 1) and (1, 1) is not present among values (*E*_*ip*_, *E*_*iq*_) so there does not exist a conflict in *E* involving columns *p* and *q*. As this holds for an arbitrary pair of mutations *M*_*p*_ and *M*_*q*_ we can conclude that *E* is conflict-free matrix.

The observation that *E* is conflict-free directly implies that

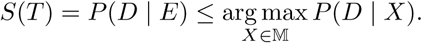

where *P* (*D* | *X*) is computed analogously as *P* (*D* | *E*) and using (1).

Since *T* is an arbitrary tree, we have

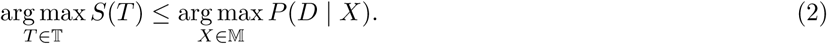

Assume now that *X* is a conflict-free matrix. Define that column *p contains* column *q* if *X*_*ip*_ ≥ *X*_*iq*_ for all rows *i*. Also, define that columns *p* and *q* are *disjoint* if there does not exist a row *i* such that (*X*_*ip*_, *X*_*iq*_) = (1, 1). It is straightforward to prove that for each two non-zero columns *p* and *q* of conflict-free matrix, either one of them contains the other or they are disjoint. Consider the rows of *X* as single-cells *C*_1_, *C*_2_, …, *C*_*n*_ and its columns as mutations *M*_1_, *M*_2_, …, *M*_*m*_ with mutation *M*_*p*_ present in cell *C*_*i*_ if and only if *X*_*ip*_ = 1. Since *X* is a binary matrix such that for each two of its non-zero columns one of them contains the other or they are disjoint, there exists a phylogenetic tree such that: (i) the set of mutations assigned to the edges of the tree equals the set {*M*_1_, *M*_2_, …, *M*_*m*_} (ii) each of the single-cells *C*_1_, *C*_2_, …, *C*_*n*_ represents exactly one leaf of the tree (iii) the set of mutations occurring on the path from the root of the tree to the cell *C*_*i*_ equals to the set of mutations present in *C*_*i*_ (this set is given by the vector [*X*_*i*1_, *X*_*i*2_, …, *X*_*im*_]). The last statement represents one of the classic results in tumor phylogenetics and its proof can be found in (Gusfield, 1991).

By detaching single-cells from the phylogenetic tree implied by *X*, we obtain a mutation tree from 𝕋. In other words, for each conflict-free matrix *X* we have corresponding mutation tree *T* such that each row of the matrix is a genotype of some node of the tree. Therefore *P* (*D* | *X*) ≤ *S*(*T*), which implies

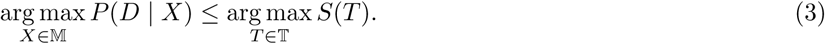

From (2) and (3) we conclude that

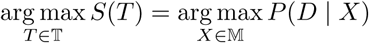

hence we can perform a search for the conflict-free matrix *X* for which *P* (*D* | *X*) is maximized and then obtain tree *T* from *X*. We also have a guarantee that, if we find an optimal matrix *X* then the related tree will be among the maximum likelihood trees (note that there might exist multiple such trees/matrices).

### 3.5 Exact integer linear programming formulation for finding maximum likelihood conflict-free matrix

In this subsection, we provide an ILP formulation to search for the conflict-free matrix *X* for which *P* (*D* | *X*) is maximized. Note that, in order to better distinguish the two tree search strategies, we opt to keep using *X* in our notation instead of using *E*.

Matrix *X* can be defined as a set of binary variables *X*_*ij*_, where *i* ∈ {1, 2, …, *n*} and *j* ∈ {1, 2, …, *m*}. First, we have to ensure that variables *X*_*ij*_ form a conflict-free matrix. In order to achieve this, we use formulation from (Gusfield et al., 2007) and introduce a set of binary variables *B*_*p,q,a,b*_, defined for each pair of columns *p* and *q* and each (*a, b*) ∈ {(0, 1), (1, 0), (1, 1)}. Our aim is that these variables satisfy the following: *B*_*p,q,a,b*_ = 1 if there exists a row *i* such that *X*_*ip*_ = *a, X*_*iq*_ = *b*. To enforce that this holds we introduce the following constraints

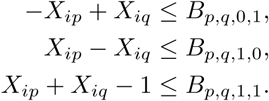

Finally, to enforce that there does not exist any conflict involving columns *p* and *q*, it is now sufficient to add the following constraint

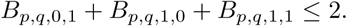

It is trivial to verify that for any conflict-free matrix *X* ∈ 𝕄 it is possible to set values of variables *B* so that all of these constraints are satisfied, hence by adding the constraints no matrix from M is excluded from the set of potential solutions.

After we have enforced that *X* is a conflict-free matrix, observe that the scoring scheme presented in (1) can be rewritten in its equivalent form

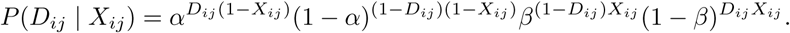

Next, note that

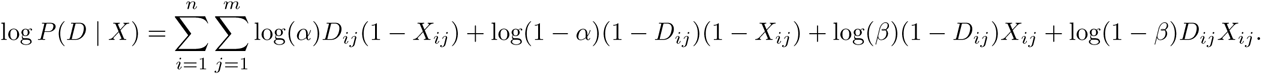

Since *D* is known, the last expression is a linear combination of unknown integer variables *X*_*ij*_. As maximizing *P* (*D* | *X*) is equivalent to maximizing log *P* (*D* | *X*), we set our objective to maximize log *P* (*D* | *X*). Clearly, we are now left with solving an instance of ILP presented above, which can be routinely done by the use of the existing ILP solvers.

## 4 The existing methods for tree inference based on the search in the space of binary matrices

The potential of the use of search in the space of binary matrices has gained a lot of attention in the past two years and several methods based on this tree-inference strategy have been proposed. In this section, we summarize five such methods and in the next section, we provide some theoretical background for another recent application.

### 4.1 PhISCS

The ILP formulation for reconstructing tumor phylogenies from single-cell data presented above is used in the implementation of PhISCS (Malikic et al., 2019b). In addition to the ILP, the equivalent Constraint Satisfaction Programming (CSP) formulation was introduced in the same work. This formulation enables the use of various, typically non-commercial, CSP solvers for solving this problem. Furthermore, the experimental results suggest that CSP implementation can be a running time-efficient alternative to the ILP implementation (without trade-off between the running time and tree reconstruction accuracy)^1^. In PhISCS, the sub-perfect phylogeny problem is also introduced and solutions based on the ILP, as well as CSP, proposed. In sub-perfect phylogeny problem, in order to account for the possible presence of violations of the infinite sites assumption, a user-specified number of mutations are allowed to be eliminated from the scoring and the reported tree, while the remaining set of mutations are assumed to satisfy the infinite sites assumption.

### 4.2 PhISCS-BnB

With the technological improvements and decrease in the cost of sequencing, we expect that many of the future SCS datasets will consist of sequencing data of hundreds or thousands of single-cells with high sequencing breadth that will enable detection of a large number of somatic single nucleotide variants. Time-efficient analysis of such datasets will require a design of more efficient computational methods that scale well to the larger inputs. Recently developed PhISCS-BnB (Sadeqi Azer et al., 2020b) is one of the methods designed to fill this gap. As demonstrated on the simulated data, PhISCS-BnB finds a solution with the optimality guarantee typically 10 to 100 times faster than PhISCS and can successfully find the optimal solution (within 24 hours) on most of the inputs as large as 300 × 300, whereas PhISCS convergence is usually limited to matrices of size 100 × 100 or smaller. In addition, the method was also compared against SCITE in terms of tree reconstruction accuracy. The obtained results suggest that, even when SCITE is given considerable running time advantage, trees reported by PhISCS-BnB on larger inputs are more similar to the ground truth simulated trees.

PhISCS-BnB is based on the branch and bound algorithm and its implementation does not require any commercial solver. Note that, in comparison to the alternative methods, it uses a simpler model in which only the presence of false negative mutation calls in the input matrix *D* is considered. Consequently, the objective function presented in Section 3 can be replaced by the objective where the goal is to minimize the sum of variables *X*_*ij*_ over the pairs (*i, j*) for which *D*_*ij*_ = 0 (i.e., minimize the number of false negatives). Since the existence of false positive mutation calls in the input is currently inevitable in most of the real applications, the robustness of the method to the presence of this type of noise was also assessed. While the existence of false positives in the input can in some cases be challenging for convergence to the optimal solution or proving that solution found within a given time limit is optimal, results suggest that, in terms of the tree reconstruction accuracy, PhISCS-BnB is robust to the presence of low levels of false positives (*α* ∼ 0.0003) as typically observed in SCS data.

### 4.3 scVILP

Due to non-uniform sequencing coverage and amplification biases inherent to single-cell sequencing, the observed total coverage and variant allele frequencies across sites and cells usually take a wide range of values. Depending on these values, we might adjust our confidence in mutational presence/absence. For example, observing 10 variant reads out of 22 total reads spanning a given genomic locus gives higher confidence in mutational presence than observing 2 variant out of 10 total reads. These differences are not well reflected in noisy genotype matrix *D* which does not contain the exact information about read counts. In scVILP (Edrisi et al., 2019), the scoring scheme presented in (1) is extended so that the read counts are directly factored in the likelihood. The input to this method consists of a matrix *H*, where *H*_*ij*_ = (*r*_*ij*_, *v*_*ij*_) represents the number of reference and variant reads observed at putative mutation loci *j* in cell *i*. Similarly, as in PhISCS, the likelihood function is a linear combination of unknown states *X*_*ij*_ with coefficients depending on read counts and a few other given constants. The search for the optimal matrix *X* is then performed in the space of conflict-free matrices analogously as described in Section 3.5.

In order to improve the running time performance, the authors also propose a divide-and-conquer approach where the matrix *H* is first split into smaller matrices *H*^(1)^, *H*^(2)^, …, *H*^(*k*)^ formed by combining subsets of columns of *H*. Then for each matrix *H*^(*i*)^ an instance of ILP is solved taking *H*^(*i*)^ as the input. Let *X*^(1)^, *X*^(2)^, …, *X*^(*k*)^ denote the optimal genotype matrices reported by these ILPs. Matrices *X*^(*i*)^ are combined into a matrix 𝒳, which obviously does not necessarily represent a conflict-free matrix. In order to eliminate conflicts in 𝒳 two steps approach is taken. First, the smallest set 𝒮 of columns needed to be removed from 𝒳 in order to obtain a conflict-free matrix is identified. In the second step, corrections of entries of 𝒳 are performed with the aim of eliminating conflicts. Importantly, only corrections allowed are these in columns from 𝒮. This usually largely reduces the size of the resulting conflict-elimination problem since the size of 𝒮 is expected to be small. In summary, after these corrections in 𝒳 are made, it is returned as a solution. This solution is not necessarily having likelihood as high as the likelihood of the optimal matrix *X*, but solving all these smaller ILP instances separately typically takes less time than solving a single instance that takes the entire *H* as the input and returns *X*.

### 4.4 SPhyR

In the k-Dollo evolutionary model used in SPhyR (El-Kebir, 2018), any SNV is gained only once but can be lost up to *k* times, where *k* is a given non-negative integer constant. Therefore an arbitrary mutation *j* can be either present at a given node or is in one of the up to *k* + 1 distinct states of mutational absence. The first of these states corresponds to the mutational absence prior to the gain of SNV, whereas each of the remaining states corresponds to mutational absence due to SNV loss (note that distinct losses can only occur on different branches of the tree). To encode the unknown true state of a mutation *j* in a cell *i* and differentiate between distinct causes for mutational absence, *k* + 2 binary variables *a*_*i,j,l*_ are introduced. The variable *a*_*i,j*,1_ is set to 1 if and only if cell *i* originates from a node that harbors mutation *j* and variable *a*_*i,j*,0_ is set to 0 if and only if cell *i* originates from a node that does not harbor mutation *j* and the absence of the mutation in this node is not due to the loss of previously gained mutation (i.e., the node is not among the descendants of the node where mutation *j* occurs). For *l* ≥ 2, the variable *a*_*i,j,l*_ is set to 1 if and only if mutation *j* is absent in cell *i* due to (*l* − 1)-st mutational loss. Combining variables *a*_*i,j,l*_ for all pairs of cells and mutations result in an unknown binary matrix *A*. Analogous to the rows of the matrix *X* introduced in Section 3, rows of *A* (indirectly) define genotypes of nodes of origin of the related cells. However, as mutational losses are allowed in this model, in order to ensure that there exists an evolutionary tree with genotypes of its nodes matching genotypes implied by rows of *A*, there are now in total (*k* + 1)^4^ + 2*k*^2^(*k* + 1)^2^ + *k*^4^ matrices of size 3 × 2 that form “conflicts” and are not allowed as submatrices of *A* (here, submatrix is any 3 × 2 matrix consisting of elements at the intersection of a pair of columns and triplet of rows of *A*). Ensuring that there does not exist any such conflict can also be achieved by the set of ILP constraints and we refer to (El-Kebir, 2018) for full details. Note that the implementation described in (*El* − *Kebir*, 2018) is based on a time-efficient heuristic. Motivated by the clonal theory of cancer evolution (Nowell, 1976; Kuipers et al., 2017), the model also allows clustering of cells and mutations, which can further improve the running time. It is assumed that all cells clustered together originate from the same node of the evolutionary tree and therefore have equal genotypes (analogous applies to the mutations).

### 4.5 gpps

The last combinatorial method that we discuss is gpps (Ciccolella et al., 2018), which is also based on the use of k-Dollo evolutionary model. From the methodological point of view, gpps is based on the elegant extension of the methodology described in Section 3. In order to allow *k* mutational losses, instead of *X*_*ij*_, a set of *k* + 1 variables *Y*_*i,j*,1_, *Y*_*i,j*,2_, …, *Y*_*i,j,k*+1_ is introduced. Intuitively, *Y*_*i,j*,1_ = 1 represents the existence of a gain of mutation *j* on the path from the tree root to the node of origin of cell *i*, whereas *Y*_*i,j,l*_ = 1, for *l* ≥ 2, represents the existence of loss of mutation *j* on such path. To ensure that the number of losses does not exceed the number of gains, the following constraint is added

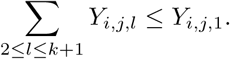

The true status (i.e., presence or absence) of mutation *j* in cell *i*, previously denoted as *X*_*ij*_, is now given by the following formula

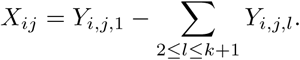

In addition to the inequality constraints presented above that limit the number of mutational losses, the matrix *Y* with *i*-th row equal to

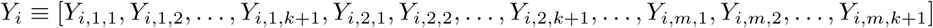

must be a conflict-free. This can be achieved by introducing a set of binary variables *B* and adding linear constraints analogously as described in Section 3. It can be shown that the above constraints are sufficient to ensure that there exists k-Dollo phylogeny consistent with *X*, i.e., genotypes of leaves (single-cells) of the phylogeny match genotypes given by rows of the matrix *X*.

As running the ILP solver until the optimal solution is found can be prohibitively slow, gpps also offers an option of running the solver for a given user-specified period of time and then running an additional algorithm based on the Hill Climbing search. The main goal of this algorithm is finding more likely phylogeny than the possibly suboptimal one reported by the ILP solver, which is used as the starting point of the Hill Climbing search.

## 5 Fast inference of the existence of divergent subclones in a given tumor via deep learning

Recently, in (Sadeqi Azer et al., 2020a) it was demonstrated that deep learning can be successfully employed in studying tumor evolution. Several distinct problems were considered and solutions to each of them involve relations between the conflict-free matrices and trees of tumor evolution. In this section, we focus on one of these problems, namely the problem of inferring whether a given tumor contains divergent subclones evolving on separate tree branches (i.e., whether the most-likely mutation tree of the tumor has a non-linear topology). For this purpose, a special subclass of binary matrices, named *staircase matrices* (see the definition below) is considered. Here, we establish some important dependencies between staircase matrices and topology of the tree of the tumor evolution. These are extensively used in (Sadeqi Azer et al., 2020a) to develop a fast deep learning based approach for discriminating between linear and non-linear tree topologies. Interestingly, the proposed solution achieves a speed-up of 100 times or more over the available alternatives.

**Definition:** A binary matrix *S*_*n*×*m*_ satisfies the *staircase* property if *S*_*ij*_ ≤ *S*_(*i*+1)*j*_ for all pairs of integers (*i, j*) such that 1 ≤ *i* ≤ *n* − 1 and 1 ≤ *j* ≤ *m*. ^2^ Any matrix satisfying the staircase property is also called *a staircase matrix*.

### Lemma 1

*If T is a mutation tree with linear topology and A is a noise-free single-cell genotype matrix, where single-cells originate from the nodes of T, then the rows of A can be reordered so that the resulting matrix satisfies the staircase property. On the other hand, if rows of a binary matrix A can be reordered so that the newly obtained matrix satisfies the staircase property, then the matrix A is conflict-free and implies a linear topology*.

**Proof:** For simplicity of notation, for any genotype matrix mentioned in this proof, the terms “row *i*”/”cell *i*” and “column *j*”/”mutation *j*” stand for the cell corresponding to the row *i* and the mutation corresponding to the column *j* of the matrix, respectively.

First, we prove that rows of a noise-free single-cell genotype matrix, where single-cells were sampled from a tree with linear topology, can be reordered so that the resulting matrix satisfies the staircase property. We first illustrate this on a simple example shown in Figure 2. Now, consider a general case and assume that we have sampled cells *C*_1_, *C*_2_, …*C*_*n*_ from a mutation tree having linear topology and the set of mutations *M*_1_, *M*_2_, …, *M*_*m*_. We sort rows of the single-cell genotype matrix such that the following is satisfied in the sorted matrix: for each *i* ≤ *n* − 1, if rows *i* and *i* + 1 respectively represent genotypes of cells *C*_*a*_ and *C*_*b*_, then the node of origin of *C*_*a*_ is either the same as the node of origin of *C*_*b*_ or is its ancestor. In other words, we sort rows in the genotype matrix from row 1 to row *n* in the same order as related cells are attached to the nodes of the mutation tree when the tree is traversed from the root towards the leaf. Let *S* denote the matrix obtained after this sorting. Our goal is to prove that *S* satisfies the staircase property, which is equivalent to proving that (*S*_*ij*_, *S*_(*i*+1)*j*_) ≠ (1, 0) for each pair of indices (*i, j*) such that 1 ≤ *i* ≤ *n* − 1 and 1 ≤ *j* ≤ *m*. In other words, we need to prove that in matrix *S* we do not have any case where mutation *j* is present in the cell *i*, but absent in the cell *i* + 1. This is true due to the construction of *S*. Namely, cell *i* originates either from the same or ancestral node of the node of origin of cell *i* + 1, hence if mutation *j* is present in the cell *i* then it must also be present in the cell *i* + 1. In other words, *S*_*ij*_ = 1 ⇒ *S*_(*i*+1)*j*_ = 1 so (*S*_*ij*_, *S*_(*i*+1)*j*_) can not be equal to (1, 0).

In the second part of the proof, assume that a binary matrix *A* can be transformed, by permuting its rows, into a matrix *S* which satisfies the staircase property. Assume also for the moment that no column in *A* consists entirely of zeros. We prove that *A* is conflict-free and that the topology of the tree implied by *A* is linear. Note that changing the order of rows in the matrix does not introduce nor resolve conflicts so *A* is conflict-free if and only if *S* is conflict-free. Similarly, the topology of the tree implied by *A* is linear if and only if the topology of the tree implied by *S* is linear. Therefore it suffices to prove that *S* is conflict-free and that the topology of the tree implied by *S* is linear. To prove that *S* is conflict-free it is sufficient to observe that for each pair of columns *p* and *q* it is not possible to find two distinct rows *i* and *j* such that (*S*_*ip*_, *S*_*iq*_) = (1, 0) and (*S*_*jp*_, *S*_*jq*_) = (0, 1). Namely, if *i < j* then we can not have *S*_*ip*_ = 1 and *S*_*jp*_ = 0 and, if *j < i*, then we can not have *S*_*jq*_ = 1 and *S*_*iq*_ = 0. Now, since *S* is conflict-free, based on the results presented in Section 3, there exists a mutation tree of tumor evolution such that the set of mutations placed at its edges corresponds to the set of columns of *S* and each row of *S*, if considered as a binary genotype, matches genotype associated with one of the nodes of the tree. Assume that topology of this tree is not linear. Then there exist mutations *p* and *q* that belong to different branches of the tree. Recall our assumption that each column of *A* (equivalently of *S*) has at least one non-zero entry. This implies that each mutation is present in at least one single-cell. Consequently, there is at least one single-cell, denote it as *i*, sampled from the subtree below the edge labeled by *p*.

Analogously, there is at least one single-cell, denote it as *j*, sampled from the subtree below the edge labeled by *q*. The cells *i* and *j* are obviously originating from different branches of the tree hence they are distinct and *i* does not harbor *q*, whereas *j* does not harbor *p*. But then we have found two distinct cells, *i* and *j*, such that (*S*_*ip*_, *S*_*iq*_) = (1, 0) and (*S*_*jp*_, *S*_*jq*_) = (0, 1), which is impossible as we have already shown above. The observed contradiction completes the proof that the tree implied by *S* is linear.

The above proof was done under the assumption that the matrix *A* does not contain a column consisting entirely of zeros. If such columns exist, then we recommend filtering them from *A*. Namely, according to *A*, such mutations are absent in all single-cells and therefore they are non-informative in tree reconstruction. Furthermore, in practice, the presence of such columns is very likely due to the false-positive mutation calls in the noisy single-cell matrix from which *A* was obtained. But if we still insist on the mutation tree which contains these mutations, we can first form a linear tree on the non-zero columns of *A* and then extend it, starting at the leaf, by adding mutations that correspond to all-zeros columns. The resulting tree is still implied by matrix *A* and has a linear topology. However, note that in this case some other topologies might describe the tumor’s evolutionary history equally well as the placement of mutations associated with all-zeros columns is uncertain.

## Footnotes

1 See also (Brown et al., 2020) for additional comparisons of ILP and CSP in solving hard problems in Computational and Systems Biology.

2 We use the term *staircase matrix* because as soon as rows of *S* can be reordered so that the inequalities *S*_*ij*_ ≤ *S*_(*i*+1)*j*_ are satisfied, we can then also sort columns of *S* from left to right in ascending order of the number of ones that they contain and color the entries equal to 1 in order to obtain the staircase-like structure. An example of such structure can be obtained by swapping the first two columns in the right matrix in Figure 2.

